# Regulatory T cell expansion by a highly CD25-dependent IL-2 mutein arrests ongoing autoimmunity

**DOI:** 10.1101/862789

**Authors:** Liliane Khoryati, Minh Nguyet Pham, McKenna Sherve, Swarnima Kumari, Kevin Cook, Josh Pearson, Marika Bogdani, Daniel J. Campbell, Marc A. Gavin

## Abstract

Interleukin-2 (IL-2) controls the homeostasis and function of regulatory T cells (Tregs) and defects in the IL-2 pathway contribute to multiple autoimmune diseases. Although recombinant IL-2 therapy has been efficacious in certain inflammatory conditions, the capacity for IL-2 to also activate inflammatory effector responses highlights the need for IL-2-based therapeutics with improved Treg-specificity. From a panel of rationally designed IL-2 variants, we identified IL-2 muteins with reduced potency and enhanced Treg-selectivity due to increased dependence on the IL-2-receptor component CD25. As an Fc-fused homodimer, the optimal Fc.IL-2 mutein induced selective Treg enrichment and reduced agonism of effector cells across a wide dose range. Furthermore, despite being a weaker agonist, overall Treg growth was greater and more sustained due to reduced receptor-mediated clearance of the Fc.IL-2 mutein compared to Fc-fused wild-type IL-2. Preferential Treg enrichment was also observed in the presence of activated pathogenic T cells in the autoimmune target organ, despite a loss of Treg-selectivity in an IL-2R-proximal response. These features allowed for extended resolution of spontaneous autoimmunity using infrequent dosing schedules. Thus, IL-2 muteins enable efficient, flexible, and targeted control of the autoimmune response.

**One Sentence Summary:** A CD25-dependent IL-2 mutein selectively expands regulatory T cells and provides potent and targeted control of autoimmunity.

## Introduction

Interleukin-2 (IL-2) is a polyfunctional cytokine that plays a central role in both the maintenance of normal immune homeostasis as well as the amplification and regulation of antigen-specific immune responses (*1, 2*). A major factor defining the systemic outcome of IL-2 signaling is whether the dominating IL-2-responsive cell populations are pro-inflammatory lymphocytes or anti-inflammatory regulatory T cells (Tregs) expressing the transcription factor forkhead box P3 (Foxp3). These differential cellular responses are dictated by the hierarchy of the IL-2 receptor (IL-2R) expression levels among these lymphocyte subsets. IL-2R is expressed either as an intermediate-affinity dimer, composed of IL-2Rβ (CD122) and the common cytokine receptor gamma chain (CD132), or a high-affinity trimer that includes IL-2Rα (CD25). CD25 has no signaling capacity but functions by binding IL-2 and presenting it to the CD122/CD132 signaling complex (*3*). Constitutive CD25 expression is largely limited to Foxp3^+^ Tregs (*4–6*), making them highly responsive to limiting amounts of IL-2. In contrast, CD122 and CD132 are found on nearly all T cells and are most highly expressed by memory CD8^+^ T cells and NK cells (*7*). Moreover, CD25 is induced on conventional CD4^+^ and CD8^+^ T cells after TCR stimulation (*8–10*), rendering them more competitive with Tregs for IL-2 access. Thus, in addition to its key anti-inflammatory functions through Treg cells, IL-2 can promote inflammation via activation of effector T cells and NK cells. Nevertheless, Treg typically express the highest levels of CD25 in inflammatory conditions due to a Treg-intrinsic positive feedback loop driven by Foxp3 enhancement of *IL2RA* transcription and IL-2R/signal transducer and activator of transcription 5 (STAT5) enhancement of *FOXP3* transcription (*11*–*14*).

The dual pro- and anti-inflammatory functions of IL-2 make it an attractive candidate for immunotherapy, and several approaches have been developed to direct IL-2 to different forms of the IL-2R to selectively activate specific target cell populations. In cancer, IL-2 can be used to stimulate CD25^-/lo^ NK cells and effector T cells to boost anti-tumoral immunity. While recombinant IL-2 (aldesleukin) was approved for cancer immunotherapy in 1992, the high doses required for tumor regression result in only limited efficacy and are associated with the drawbacks of significant Treg activation and severe side effects such as vascular leak syndrome leading to multi-organ dysfunction (*15, 16*). More recently, mutated IL-2 fusion proteins, IL-2 mutants, or *de novo* proteins have been engineered to overcome these limitations by deleting the vasopermeability activity of IL-2 (*17–19*) or directing IL-2 binding towards the CD122/CD132 heterodimer and away from CD25 (*20, 21*). Conversely, for autoimmune and inflammatory diseases, the objective has been to leverage IL-2 to enhance Treg numbers and function while minimizing activation of pathogenic effector cells. Here, IL-2 has been tested at the lowest biologically active doses with carefully optimized dosing regimens (*22–24*). However, balancing the efficacy and the safety of IL-2 therapy in these settings has remained a major concern, providing further support for the development of IL-2-based therapeutics with improved Treg-selectivity.

IL-2 variants (IL-2 muteins) with decreased CD122 affinity—dimerized through Fc or IgG fusion (Fc.IL-2 mutein or IgG.IL-2 mutein, respectively) for increased half-life and enhanced CD25 avidity—represent one approach for increasing Treg-selectivity. Recently, a human IgG.IL-2 mutein was developed that selectively activated and expanded functional Tregs in humanized mice during xenogeneic graft-versus-host disease (GVHD) (*25*), and an Fc.IL-2 mutein has advanced to clinical trials in GVHD (NCT03422627), systemic lupus erythematosus (SLE) (NCT03451422) and rheumatoid arthritis (RA) (NCT03410056). Despite these advances, detailed mechanistic studies evaluating the therapeutic effect of IL-2 muteins on Tregs, effector T cells and NK cells in the secondary lymphoid organs and inflamed target tissues are lacking. Evaluating IL-2 muteins in mouse disease models is necessary to define their mechanism of action *in vivo*, and can be used to help develop and guide therapeutically efficacious dosing strategies. To this end, we generated and screened a panel of 28 candidate muteins, and identified a highly Treg-selective molecule with potent Treg-enhancing properties. Through *in vitro* and *in vivo* studies, we explored the functional mechanisms of enhanced mutein activity compared to wild-type (WT) IL-2 and other previously reported Treg-boosting strategies. Finally, using a pre-clinical mouse model of type 1 diabetes (T1D) we demonstrated potent and lasting efficacy of IL-2 mutein treatment in arresting ongoing autoimmunity, with significantly greater dosing flexibility compared to WT IL-2.

## Results

Reducing IL-2 binding to CD122 can increase CD25 dependency (*26*), and thereby improve Treg-selectivity in therapeutic settings. To this end, we generated constructs for twenty-eight mouse IL-2 variants (IL-2 muteins, table S1), mutating conserved residues with known functional importance at the CD122 contact interface (*27–29*). To improve pharmacokinetics and tissue distribution, the IL-2 muteins were fused to an IgG2a Fc domain which contained an N297G mutation to abolish effector function (*50*) (hereafter referred to as Fc.WT and Fc.muteins). We first expressed the Fc.IL-2 constructs in HEK 293T/17 cells and screened the supernatants for activity by measuring phosphorylation of STAT5 in mouse splenic Tregs. Fc.WT showed comparable activity to mouse recombinant IL-2 in stimulating STAT5 phosphorylation in Tregs, Foxp3^−^CD4^+^ T cells, CD8^+^ T cells, and NK cells, indicating that the fusion of murine IgG2a Fc domain to IL-2 did not affect its bioactivity or IL-2R-selectivity (fig. S1). The Fc.muteins ranged from full activity on Tregs comparable to Fc.WT to complete functional abrogation (fig. S2). Since weaker mutein activity (indicating diminished IL-2/CD122 interaction) should translate into enhanced CD25-dependence and improved Treg-selectivity, we selected four mutein candidates exhibiting intermediate (Fc.Mut24 and Fc.Mut12) to weak/negligible (Fc.Mut25 and Fc.Mut26) activity for further evaluation. When titrated on freshly isolated splenocytes, Fc.Mut24 (EC50=11.8pM), Fc.Mut12 (EC50=162.3pM) and Fc.Mut25 (EC50=1046pM) showed reduced signaling activity on Treg cells compared to Fc.WT (EC50=0.058pM), whereas Fc.Mut26 was functionally inactive (Fig. 1, A and B). Unlike Fc.WT, the Fc.muteins did not stimulate Foxp3^−^CD4^+^ T cells, CD8^+^ T cells or NK cells even at high doses (Fig 1, A and B). Although Fc.WT stimulated nearly all Treg cells, STAT5 phosphorylation in response to the Fc.muteins was limited to CD25^hi^ Treg cells; and decreased potency of the Fc.muteins correlated with enhanced CD25 dependence for signaling (Fig 1, A to C). Using CD25-expressing HEK cells, we found that the Fc.muteins bound CD25 with a comparable affinity to Fc.WT (fig. S3), indicating that their reduced signaling potencies were due to impaired interaction with CD122. To assess the Treg-selectivity of the Fc.muteins under conditions more closely resembling inflamed tissues, we used splenocytes that were preactivated with anti-CD3, which results in elevated CD25 expression and enhanced IL-2 responsiveness of both Foxp3^−^CD4^+^ and CD8^+^ T cells. Although as expected we observed increased Fc.mutein activity on activated effector T cells, the Fc.muteins retained enhanced Treg-selectivity and CD25-dependence compared to Fc.WT across a wide dose range (fig. S4).

**Fig. 1.**
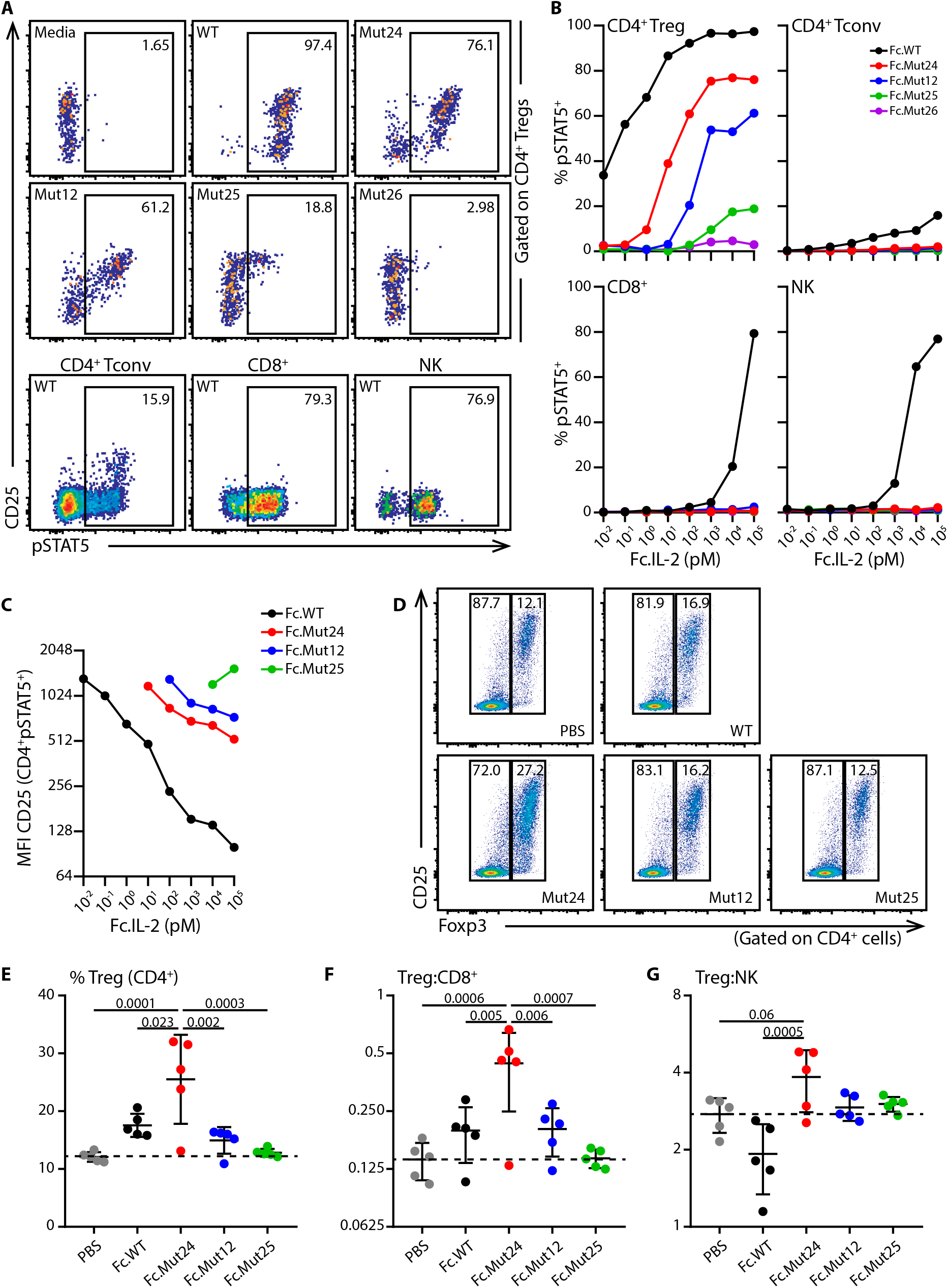
Fc.muteins show a weak signaling potency but a high Treg-selectivity. (**A to C**) B6 Splenocytes were stimulated with the indicated purified Fc.IL-2 fusion proteins. (**A**) Representative flow cytometric analyses for the 100nM dose. (**B**) pSTAT5 dose-response curves for the indicated populations. (**C**) CD25 median fluorescence intensity on CD4^+^pSTAT5^+^ cells. Data are representative of at least two independent experiments. (**D to G**) B6 mice were treated on day 0 with a single dose of PBS or the indicated Fc.IL-2 (8μg), and inguinal lymph nodes were harvested four days later. (**D**) Representative flow cytometric analyses of Treg frequency. (**E**) Frequencies of Treg and ratios of (**F**) Treg:CD8^+^ and (**G**) Treg:NK are summarized. Data shown as mean ± SD, significance determined by one-way ANOVA followed by Tukey post-test. Representative of at least two independent experiments with n=5 mice/group.

To extend these findings *in vivo*, we compared responses of IL-2 sensitive lymphocytes in the lymph nodes (LN) of B6 mice after a single treatment (8μg) with Fc.WT, Fc.Mut12, Fc.Mut24 and Fc.Mut25. Surprisingly, despite its weaker activity *in vitro*, Fc.Mut24 was more effective than Fc.WT at expanding the Treg population relative to conventional Foxp3^−^CD4^+^ T cells, CD8^+^ T cells and NK cells *in vivo*, whereas Fc.Mut12 showed weaker activity and Fc.Mut25 had no effect (Fig. 1, D to G). Based on these results we selected Fc.Mut24 as the optimal candidate for mechanistic studies of Treg-enrichment based therapy in autoimmunity.

The narrow dose-range that must be used to avoid effector cell activation is a barrier to the implementation of IL-2-based therapies in autoimmune diseases. To determine if the increased CD25-dependency of Fc.Mut24 would enable a wider-dose range while maintaining Treg-selectivity, we assessed Treg and effector lymphocyte activation in response to increasing doses of Fc.WT or Fc.Mut24. As indicated by induction of the proliferation marker Ki-67, both Fc.WT and Fc.Mut24 activated Tregs in a dose-dependent manner. However, whereas Fc.WT also activated Foxp3^−^CD4^+^ T cells, CD8^+^ T cells and NK cells, Fc.Mut24 did not activate any effector T cells and only stimulated NK cells at the highest doses tested (Fig. 2A). Consequently, there was a robust dose-dependent increase in the frequency of Tregs compared to all of the effector populations in Fc.Mut24-treated animals (fig. S5), with Tregs reaching more than 50% of CD4^+^ T cells at the highest dose. In contrast, Treg frequencies in mice given Fc.WT plateaued at less than 30% (Fig. 2, B to E). Stimulation of NK cells by Fc.Mut24 *in vivo* is curious given their lack of CD25 expression and the inability of Fc.Mut24 to induce STAT5 phosphorylation in NK cells *in vitro*. However, NK cells express extremely high levels of CD122, and trans-presentation of IL-2 by CD25-expressing cells to CD122^hi^ cells has been reported (*31*). We therefore speculate that the activation of NK cells observed with Fc.Mut24 may involve trans-presentation by soluble CD25 or CD25-expressing cells, and this NK response may represent the limit of Treg specificity that can be achieved by selectively targeting IL-2 to CD25^+^ cells. Administration of daily low-dose human IL-2, or IL-2/JES6-1A12 immune complexes (IL-2ic) have also been used to enrich Tregs and treat autoimmunity in mice (*32*). However, Fc.Mut24 treatment drove significantly greater Treg expansion compared to either of these approaches, and in particular daily low-dose human IL-2 treatment had a very modest effect on Treg enrichment despite showing an equivalent activity to Fc.WT and murine recombinant IL-2 *in vitro* (fig. S6).

**Fig. 2.**
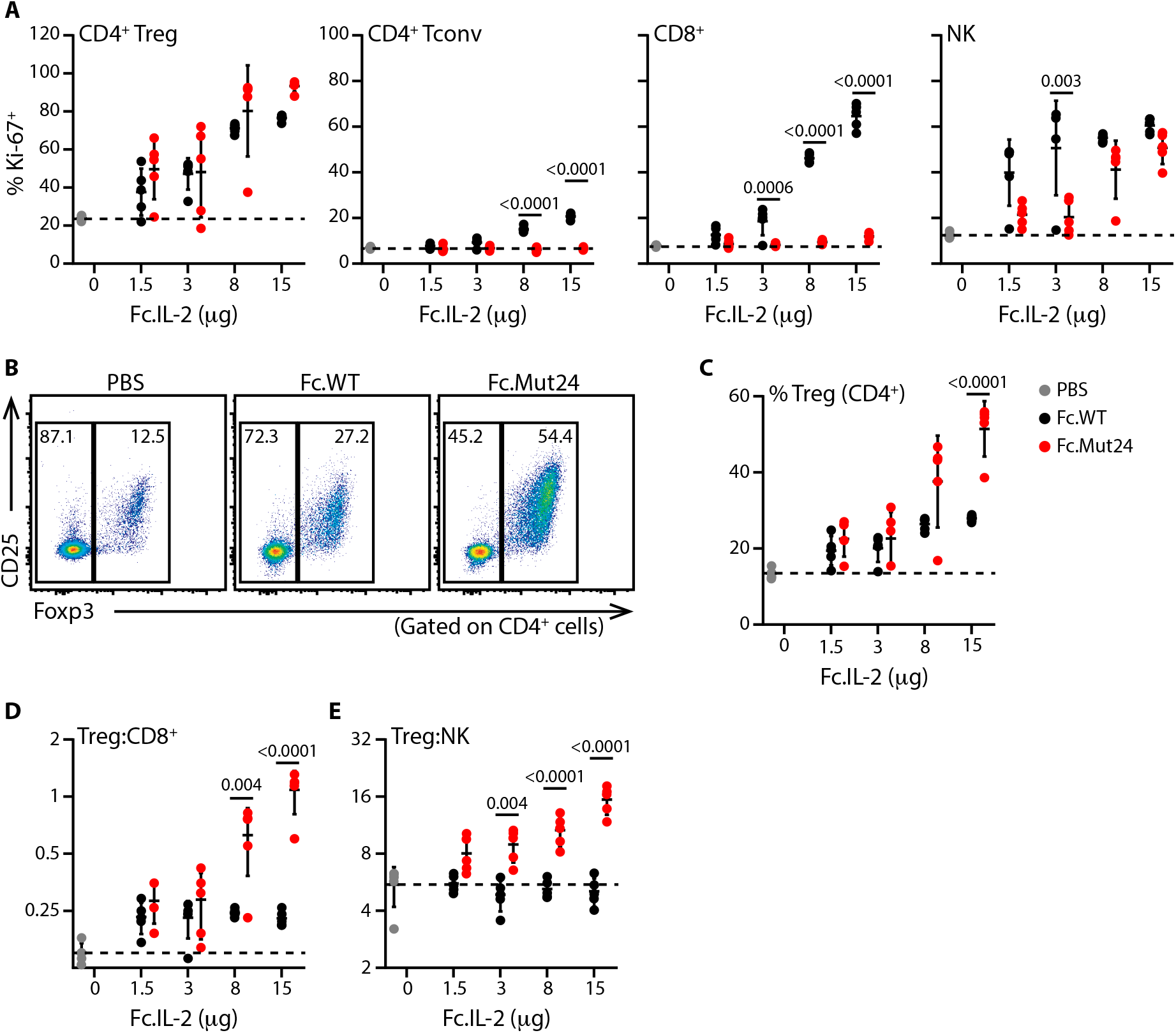
Fc.Mut24 remains Treg-selective across a wide dose range *in vivo*. B6 Mice were treated on day 0 with either PBS or Fc.IL-2 and inguinal lymph nodes were harvested four days later for analysis by flow cytometry. (**A**) Percentages of Ki-67^+^ cells in the indicated populations are plotted. (**B**) Representative analyses of Foxp3 and CD25 expression for the 15μg group. (**C**) Treg frequencies and ratios of (**D**) Treg:CD8^+^ and (**E**) Treg:NK are summarized. Data shown as mean ± SD, significance determined by two-way ANOVA followed by Tukey post-test. Representative of at least two independent experiments with n=5 mice/group.

We next examined the kinetics of Treg enrichment in response to a single high-dose (15μg) treatment with either Fc.WT or Fc.Mut24. In Fc.WT-treated mice, the percentage of Tregs in the blood peaked on day 3 and declined rapidly to near baseline by day 7. By contrast, the response to Fc.Mut24 peaked on day 4, then slowly declined but remained elevated compared to baseline for at least a week (Fig. 3, A and B). Furthermore, at all timepoints the ratios of Treg:CD8^+^ cell and Treg:NK cell were higher in Fc.Mut24-treated mice than in Fc.WT-treated ones, indicating that Treg-selectivity is maintained throughout the response (Fig. 3, C and D). At days 5 and 7 post-treatment, Treg expansion and selectivity in the LN of Fc.Mut24-treated mice vs Fc.WT-treated mice was similar to that in the blood (fig. S7). Thus, despite being a weaker agonist of IL-2R signaling *in vitro*, we observed more robust and sustained responses of Treg cells to Fc.Mut24 *in vivo*. The *in vivo* half-life of Fc.IL-2 fusion proteins is largely a function of receptor mediated consumption (*25, 33*). This led us to hypothesize that due to its weaker activity, Fc.Mut24 should exhibit reduced receptor-mediated clearance, resulting in prolonged retention on the surface of CD25^+^ cells and an extended *in vivo* half-life. To test this, splenocytes preactivated with anti-CD3/CD28 were coated with Fc.WT or Fc.Mut24, washed and cultured at 37°C for up to 24h. We then measured the amount of IL-2 fusion protein on the cell surface by staining cells for CD25 and the Fc portion of cell-associated Fc.IL-2. Whereas the amount of Fc.WT on the surface of CD4^+^CD25^hi^ cells rapidly declined, surface association of Fc.Mut24 was dramatically prolonged, and we detected substantial levels of Fc.Mut24 on the cell surface even after 24h (Fig. 3, E and F). Accordingly, the half-life of Fc.Mut24 *in vivo* was approximately three times longer than Fc.WT, and was closer to that of the functionally inactive mutein Fc.Mut26 (Fig. 3, G and H). Thus, diminished consumption via IL-2R signaling resulted in extended pharmacokinetics and sustained activity of Fc.Mut24, and this likely explains the potent Treg enrichment observed after Fc.Mut24 treatment.

**Fig. 3.**
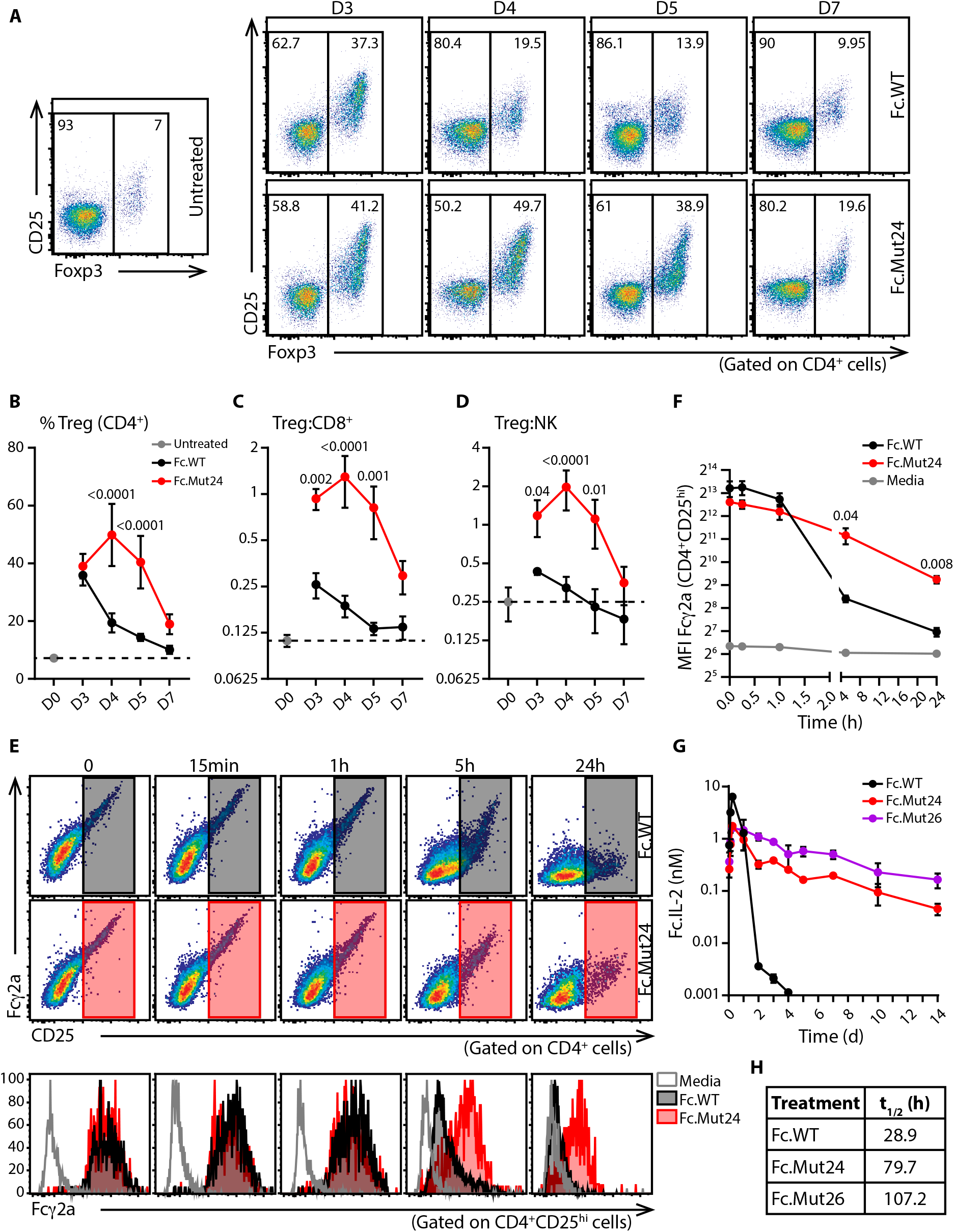
Sustained Treg enrichment *in vivo* and lower target mediated clearance of Fc.Mut24. (**A to D**) B6 Mice were treated on day 0 with Fc.WT or Fc.Mut24 (15μg) and blood was collected at different times for analysis. (**A**) Representative analyses of Foxp3 and CD25 expression. (**B**) Treg frequencies and ratios of (**C**) Treg:CD8^+^ and (**D**) Treg:NK are summarized. Data shown as mean ± SD, significance determined by two-way ANOVA followed by Tukey post-test. Representative of two independent experiments with n=4-5 mice/group. (**E and F**) Anti-CD3/CD28 activated B6 splenocytes were coated with Fc.WT or Fc.Mut24 (1nM), washed, and cultured for the indicated times. (**E**) Upper and middle: Representative staining for surface Fc.IL-2 and CD25 on CD4^+^ cells. Bottom: Analysis of Fcγ2a staining on CD4^+^CD25^hi^ cells. (**F**) Analyses of median fluorescence intensity of Fcγ2a on CD4^+^CD25^hi^ cells. Data shown as mean ± SD, significance determined by repeated-measures two-Way ANOVA with the Geisser-Greenhouse correction followed by Tukey post-test. Data representative of three biological replicates. **(G and H)** Female B6 mice were administered a single dose of Fc.IL-2 (4μg) and blood was collected at different times for pharmacokinetic analysis. **(G)** Serum levels and (**H**) *in vivo* half-life of the indicated Fc.IL-2 molecules. Data shown as mean ± SD, n=3 mice/time point.

In the non-obese diabetic (NOD) model of T1D, autoimmunity begins as early as 3 weeks of age (*34*), and is characterized by infiltration of CD25^+^ effector T cells in the pancreas (*32*). Therefore, to determine if the Treg-selectivity of Fc.Mut24 is maintained at sites of ongoing autoimmunity, we treated 9-week old female NOD mice with a single 15μg dose of Fc.WT, Fc.Mut24, or PBS, and assessed responses of different lymphocyte populations in the pancreas, pancreatic lymph node (pLN), and spleen. Unlike the Treg-selective Ki-67 responses to Fc.Mut24 observed in the pLN and spleen, Fc.Mut24 and Fc.WT induced Ki-67 expression in pancreatic Foxp3^-^CD4^+^ and CD8^+^ T cells to a similar degree (Fig. 4A and fig. S8). However, despite this apparent loss of Treg-selectivity in the target organ of the autoimmune response, we still observed dominant expansion of Tregs in all tissues of Fc.Mut24-treated NOD mice that was superior to that in Fc.WT-treated animals (Fig. 4, A to D).

**Fig. 4.**
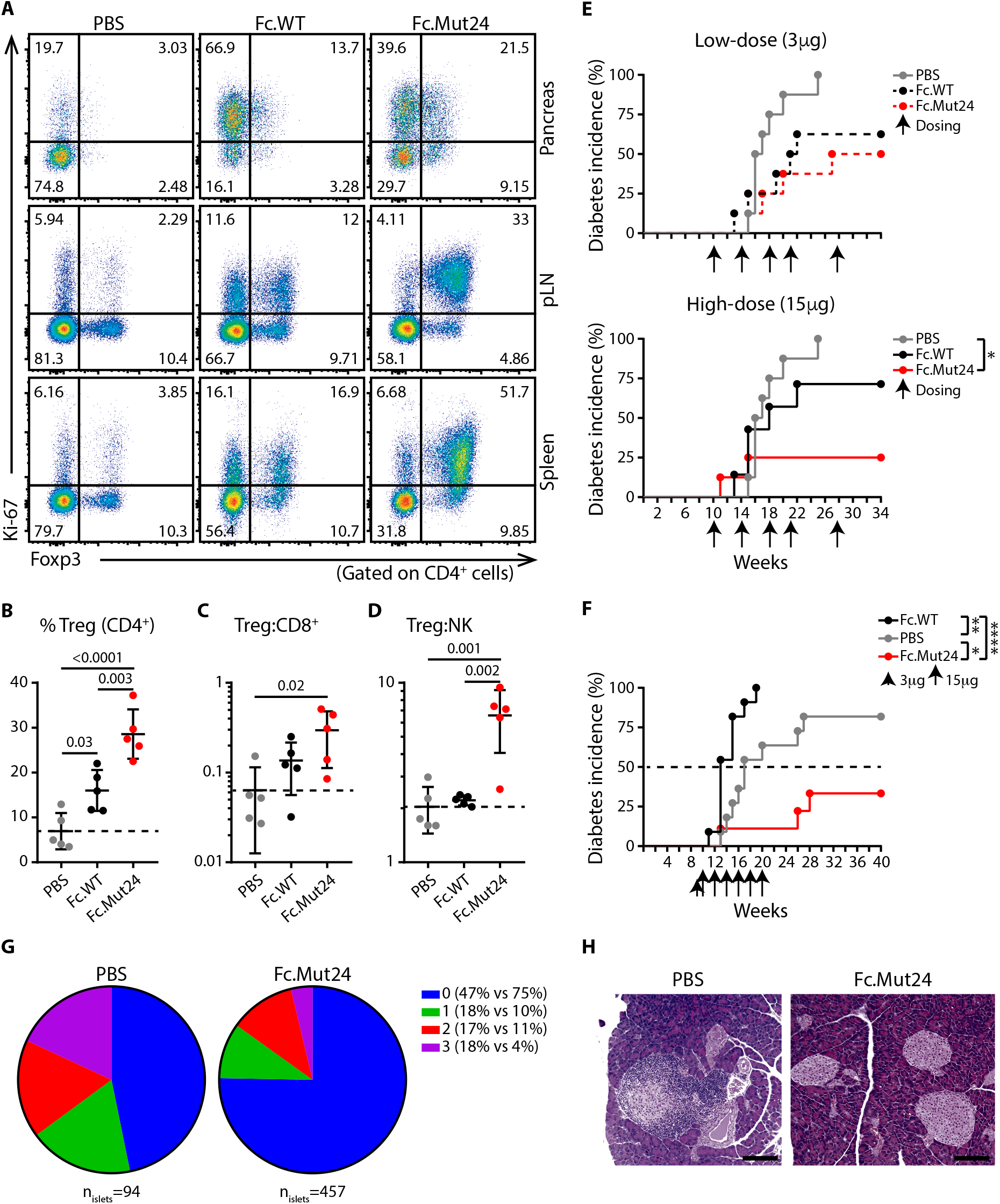
Fc.Mut24 induces selective Treg enrichment and arrests ongoing autoimmunity in NOD mice. (**A to D**) Female NOD mice were treated on day 0 with PBS, Fc.WT or Fc.Mut24 (15μg) and tissues were harvested four days later. (**A**) Representative analyses of Foxp3 and Ki-67 expression in the indicated tissues. (**B**) Treg frequencies and ratios of (**C**) Treg:CD8^+^ and (**D**) Treg:NK in the pancreas. Data shown as mean ± SD, significance determined by one-way ANOVA followed by Tukey post-test. Representative of two independent experiments with n=5 mice/group. (**E and F**) Diabetes incidence in female NOD mice treated with PBS, Fc.WT or Fc.Mut24. Arrows indicate time of dosing. Significance determined by Log-rank (Mantel-Cox) test, *p<0.05, **p<0.01, ****p<0.0001, n=7-8 in (**E**) and n=9-11 in (**F**). (**G and H**) Histological analysis of insulitis severity in disease-free mice at the end of the experiment shown in (**F**). (**G**) Percentages of islets with no infiltration (0), peri-infiltration (1), <50% infiltration (2), and >50% infiltration (3). (**H**) Representative H&E staining of a grade 3 islet from a PBS-treated mouse and three grade 0 islets from an Fc.Mut24-treated mouse (scale bar 100μm).

In the NOD T1D model, daily low-dose IL-2 treatment led to significant disease protection whereas high-dose IL-2 treatment actually accelerated disease by potentiating effector T cell and NK cell responses (*32*). Thus, this an ideal model to determine if the improved Treg-selectivity and pharmacokinetics of Fc.Mut24 translate to better therapeutic efficacy across a wide dose-range and with reduced dosing frequency. To test this, we treated NOD mice (beginning at 10 weeks of age) with either a 3μg or 15μg dose of Fc.WT or Fc.Mut24 a total 5 times over 18 weeks, and monitored blood glucose levels until 34 weeks of age. By 25 weeks, all PBS-treated control mice developed diabetes. In contrast, treatment with 3μg doses of Fc.WT or Fc.Mut24 treatment offered substantial disease protection. Whereas the efficacy of Fc.WT was somewhat reduced at the higher dose, 15μg doses of Fc.Mut24 further enhanced protection against diabetes, with 75% of the mice remaining disease-free throughout the experiment (Fig. 4E). This indicates that clinical efficacy can be achieved by Fc.Mut24 treatment across a wide dose range and with very infrequent dosing.

To further evaluate the therapeutic efficacy of Fc.Mut24, we compared Fc.WT and Fc.Mut24 in the NOD model using a dose escalation regimen in which mice were given a single 3μg dose, followed 4 days later by a 12 week course of bi-weekly 15μg treatments. Consistent with previous studies examining frequent high-dose IL-2 administration, in this regimen Fc.WT treatment markedly accelerated disease onset compared to PBS-treated controls. In contrast, Fc.Mut24 conferred a significant and durable protection against diabetes that in the majority of treated animals was sustained for at least 20 weeks following treatment cessation (Fig. 4F). Consistent with arrest of the autoimmune response, histological examination of the mice that were diabetes-free at the end of the study showed a dramatic reduction in the severity of insulitis in the Fc.Mut24-treated mice compared to the surviving PBS-treated controls (Fig. 4, G and H). Taken together, our studies in NOD mice demonstrate that Fc.Mut24 treatment results in efficient, targeted, and sustained control of autoimmunity, despite the presence of Fc.mutein-responsive pathogenic T cells at the site of autoimmune inflammation.

## Discussion

Defects in Treg cell function and alterations in the IL-2 pathway are associated with autoimmunity (*35, 36*). These findings have sparked interest in using IL-2 immunotherapy to re-establish tolerance in patients with autoimmune and inflammatory diseases. Administered at low-doses to prevent the activation of pathogenic effector cells, IL-2 therapy showed promising results of reduced disease activity and improved outcomes in patients with hepatitis C virus-induced vasculitis (*37*), graft-versus-host disease (*38–40*), alopecia areata (*41*), and systemic lupus erythematosus (*42–46*). Moreover, a recent clinical trial investigating the effects of low-dose IL-2 therapy across 11 autoimmune diseases has shown encouraging signs of efficacy using a dose of 1 MIU/injection in all patients (*47*). However, although these studies highlight the promise of IL-2-based therapies in autoimmunity, establishing a “universal” dosage that reliably induces robust Treg responses in all patients is likely to be quite challenging, as considerable heterogeneity is observed in responses to low dose IL-2 strategies (*24*). Differences in the physiopathology of autoimmune diseases may be associated with variable frequencies of CD25-high pathogenic Teff in the target tissue, thereby altering the dose required for optimal Treg selectivity. Furthermore, certain patients may carry intrinsic alterations of the IL-2 pathway that can limit the efficacy of the treatment, as in T1D (*48, 49*). To avoid potential treatment-induced toxicity and disease exacerbation by activating effector responses, the therapeutic dose-range for low-dose IL-2 remains narrow and potentially too low to achieve optimal clinical outcomes. Finally, the short *in vivo* half-life of IL-2 necessitates very frequent administration. Thus, there is a pressing need for new IL-2-based therapeutics with enhanced Treg-selectivity and improved pharmacokinetics.

We have engineered cell-selectivity into IL-2 by introducing defined point mutations at the interface with CD122. From a panel of 28 candidates, we selected the mutant Fc.Mut24 that showed increased CD25-dependency compared to Fc.WT and optimal Treg enrichment *in vivo*. Fc.Mut24 contains two amino acid substitutions at conserved CD122 contact residues, replacing neutral-polar residue N103 with a positively charged arginine (R), and hydrophobic V106 with a negatively charged aspartic acid (D). Based on IL-2 and IL-2Rβ mouse:human homologies and structures of the human IL-2/IL-2R quaternary complex, N103 in murine IL-2 (homologous to N88 in human IL-2, N88^hIL-2^) should form interactions with 4 conserved residues on murine IL-2Rβ (R42, Q71, T74, Y135), and V106 in murine IL-2 (V91^hIL-2^) should interact with 3 conserved murine IL-2Rβ residues, 2 of which also interact with N88^hIL-2^ in the human structure (R42, T74, V76) (*3, 50*). Interestingly, R41 in human IL-2Rβ (R41^hIL-2Rb^) also contacts V91^hIL-2^, and is the only residue involved in contacting either N88^hIL-2^ or V91^hIL-2^ that is not conserved in murine IL-2Rβ. Given that the analog of the positively charged R41^hIL-2Rβ^ is a hydrophobic L41 in mouse (L41^mIL-2Rβ^), we speculate that the presence of a leucine at position 41 may actually increase the strength of interaction with the hydrophobic V106 in murine IL-2, compared to R41^hIL-2Rβ^-V91^hIL-2^ interaction in human. This may explain why both the N103R and V106D mutations were required to achieve the substantially reduced signaling potency of Fc.Mut24. Indeed, the N88D and N88R mutations substantially attenuated the function of human IL-2 (*25, 26*), whereas their homologous N103R mutation in murine IL-2 only modestly affected signaling potency (Fc.Mut11 in fig. S2). Thus, the combination of the N103R and V106D mutations in Fc.Mut24 are expected to partially disrupt IL-2 binding to a specific region of IL-2Rβ.

Fc.Mut24 induced a robust expansion of Tregs in multiple locations including the blood, secondary lymphoid tissues, and non-lymphoid autoimmune target tissue of treated mice. Thus, differences in the Treg expansion following Fc.WT and Fc.Mut24 treatment are not due to differential effects on Treg recruitment to different tissue compartments, but instead reflects the short-term impact of Fc.WT on Treg enrichment compared to the extended response induced by Fc.Mut24. The *in vivo* half-life of Fc.IL-2 fusion proteins depends on the rate of IL-2R mediated clearance (*25, 33*). Consistent with this, we found that the half-life of Fc.WT, Fc.Mut24 and Fc.Mut26 inversely correlated with their signaling potencies, and that the half-life of Fc.Mut24 is almost three times longer than Fc.WT. *In vitro*, we observed a prolonged association of Fc.Mut24 with surface CD25. The slower clearance of Fc.Mut24 off the surface of CD25-expressing cells would allow for extended bioavailability compared to a rapid consumption of Fc.WT, and the improved pharmacokinetics of Fc.Mut24 likely underpin its extended activity and enhanced Treg cell enrichment compared to Fc.WT.

The increased CD25-dependency and prolonged *in vivo* half-life of Fc.Mut24 translate into an improved dynamic range of selective Treg enrichment and a greater flexibility in therapeutically efficacious dosing strategies. Furthermore, we found that chronic Treg enrichment achieved by frequent dosing was not required to provide lasting protection against autoimmunity, and that durable responses could be observed for up to 20 weeks after cessation of treatment. Durable reversal of autoimmunity occurred despite efficient Ki-67 upregulation in pancreatic effector T cells, presumably due to their high CD25 expression. This loss of Treg selectivity in inflamed tissues suggests that systemic Treg enrichment may be important for resolving organ-specific autoimmunity. Currently, human IL-2 muteins with analogous Treg-selective properties are being tested for potential use in autoimmune and inflammatory diseases (*51, 52*). A recent clinical trial treated high-risk first-degree relatives of T1D patients who tested positive for at least two diabetes-related autoantibodies with anti-CD3, and showed significant therapeutic benefit in delaying disease onset (*53*). Similarly, we started Fc.IL-2 treatment in NOD mice after initiation of autoimmunity but before development of overt hyperglycemia, and our results suggest that IL-2 mutein treatment may be also therapeutically efficacious in at-risk subjects. Thus, Fc.Mut24 is a valuable experimental tool for pre-clinical mechanistic studies, and our findings provide a strong proof-of-concept for IL-2 mutein therapy that help inform future design of clinical trials and dosing strategies. Through optimal and targeted control of the autoimmune response, the IL-2 mutein approach represents a more efficient and flexible alternative than low-dose IL-2 treatment for successful control of autoimmunity.

## Materials and Methods

### Panel design and production of Fc.IL-2 muteins

Using a site-directed mutagenesis approach on mouse IL-2, constructs for twenty-eight IL-2 muteins with point mutations introduced at the CD122 interface were generated. All muteins also contain P51T and C140A mutations, which improved manufacturability but did not affect IL-2 activity. IL-2 muteins were fused via a 4GS linker to the C-terminus of murine IgG2a Fc (N297G, deleted c-terminal K). Fc.IL-2-expressing plasmids (pcDNA3.1) were transiently transfected into HEK 293T/17 adherent cells (ATCC CRL-11268) using PEI 25K (Polysciences, Inc.) (ratio 3:1, PEI to DNA (w/w)). Three days later, supernatants were harvested for *in vitro* screening. Selected IL-2 muteins were produced and purified by the Biologics Production Facility at Fred Hutchinson Cancer Research Center (Seattle, WA) or by Olympic Protein Technologies (Seattle, WA). Purified IL-2 muteins used in *in vivo* experiments contained less than 15 EU/mL of endotoxin.

### Mice

C57BL/6J (B6) (stock number: 000664) and NOD/ShiLtJ (NOD) (stock number: 001976) mice were purchased from The Jackson Laboratory. All animal experiments were conducted in accordance with policies of the Institutional Animal Care and Use Committee of the Benaroya Research Institute, or the Amgen Institutional Animal Care and Use Committee.

### *In vitro* assays

For pSTAT5 assays, freshly isolated or anti-CD3 (1μg/mL, 145-2C11, BD Pharmingen) activated B6 splenocytes (5×10^5^ cells) were stimulated with titrated human recombinant IL-2 (Proleukin, Novartis), mouse recombinant IL-2 (eBioscience), or Fc.IL-2 for 15 min at 37°C in complete RPMI (RPMI (Hyclone) supplemented with 10% FBS (heat inactivated, VWR), L-glutamine, penicillin-streptomycin, and sodium pyruvate (Gibco)). For Fc.IL-2 association with cell surface CD25, B6 splenocytes were activated with 1μg/mL anti-CD3 and 1μg/mL anti-CD28 (37.51, BioXCell) for 24h. Cells were washed, rested for 48h, then coated with 1nM Fc.IL-2 for 15 min at RT. Coated splenocytes were thoroughly washed (3 times) to remove excess protein and incubated at 37°C for various time points. All the steps were performed in complete RPMI. For Fc.IL-2 binding to CD25 assay, mouse CD25-expressing plasmid (Sino Biological) was transiently transfected into HEK 293T/17 adherent cells using PEI 25K (ratio 3:1, PEI to DNA (w/w)). CD25-expressing HEK 293T/17 cells were coated with titrated Fc.IL-2 for 15 min at RT in DMEM (Hyclone) supplemented with 5% FBS. Coated cells were thoroughly washed (4 times) then stained for flow cytometry analysis.

### *In vivo* dosing assays

B6 mice (7-12 weeks old, females or males) and NOD mice (9-10 weeks old, females) were administered on day 0 a single dose of Fc.IL-2, IL-2 immune complex (IL-2ic; mouse recombinant IL-2 + anti-IL-2 (JES6-1A12, BioLegend); molar ratio 2:1, IL-2 to antibody), or PBS by intraperitoneal (IP) injection. Human recombinant IL-2 was administered by daily IP injection for four consecutive days (day 0 through day 3). Four days after initiation of the treatment (day 4), mice were euthanized and tissues were harvested for analysis. For the time course experiment, blood was collected from the saphenous vein of untreated mice (day 0) and on day 3, 4, 5, and 7 after dosing. Inguinal lymph nodes were harvested on day 0, 5, and 7.

### PK study

All procedures were in compliance with the Animal Welfare Act Regulations (9 CFR 3) overseen and approved by the Amgen Institutional Animal Care and Use Committee. Female C57BL/6 mice (Charles River Laboratories) received a 4 μg/mouse subcutaneous bolus dose of Fc.IL-2 in the mid-scapular region. At each time point approximately 0.06 mL of whole blood per mouse was collected via submandibular vein puncture and processed into serum. Quantitation of IL-2 muteins in mouse serum was performed using an electrochemiluminescent immunoassay with a biotinylated rat anti-mouse IL-2 (JES6-1A12, R&D Systems) as a capture reagent and a ruthenylated goat anti-mouse IgG, Fcγ subclass 2a specific (Jackson ImmunoReasearch Laboratories) as a detection reagent. Analyte serum concentrations were interpolated from a standard curve using the corresponding analyte. Noncompartmental pharmacokinetic analyses were performed for the mean of three study subjects at each time point using non GLP Watson LIMS (Thermo Fisher Scientific).

### NOD mice diabetes study

Female NOD mice were received at 6 weeks of age. Cages were changed once a week for mice younger than 10 weeks old and twice a week after that. Blood glucose levels (BGV) and body weight were measured weekly. Bayer Contour next meter and Contour next Blood glucose test strips were used for BGV monitoring. Diabetes was confirmed when BGV levels were above 250mg/dL for two consecutive days and diabetic mice were euthanized within 24h of the second measurement.

In our dosing strategy for the first study, mice received IP injections of PBS or a low (3μg) or high (15μg) dose of Fc.WT or Fc.Mut24 at 10 weeks of age, and subsequent doses were administered to all groups when a single mouse in the low-dose Fc.Mut24 group became diabetic. In this manner, the rate of disease onset for the low-dose Fc.Mut24 group was governed by the loss of immunosuppression following individual doses, and the rate of onset in the other groups would reveal if the same dosing frequency would be more or less effective at controlling disease. A total of five IP injections were administered between week 10 and 28. At week 34, all the mice that remained diabetes-free were euthanized.

A second cohort of NOD mice was treated beginning at 10 weeks of age with a short dose-escalation phase by administering the mice a low-dose (3μg) of Fc.IL-2 followed by a high-dose (15μg) four days later. Maintenance high-dose injections (15μg) were then given once every two weeks between weeks 12-20. At week 40, all the mice that remained diabetes-free were euthanized.

### Flow Cytometry

Single cell suspensions from spleen and LN were prepared by mechanic disruption. Pancreata were digested enzymatically in plain RPMI (Hyclone) supplemented with DNase I (1:200, D4513, Sigma-Aldrich) and liberase TM (1:100, 5401127001, Roche) for 30 min at 37°C under agitation. Cell suspensions were filtered through 70μm strainer and immune cells were isolated using Percoll gradient (GE Healthcare Life Sciences). For blood samples, RBCs were lysed using the 1X RBC lysis buffer (eBioscience).

Extracellular staining was performed in PBS supplemented with 0.5% BSA (Sigma-Aldrich) at 4°C. For surface mutein staining, samples were incubated first with polyclonal anti-mouse Fcγ2a PE (Jackson ImmunoResearch Laboratories). After 30 min, cells were washed and mouse IgG2a (HOPC-1, BD Pharmingen) was used for blocking for 15 min. Surface staining antibodies were then added for 30 min.

Intracellular staining was performed using the Foxp3 transcription factor staining buffer set (eBioscience). For pSTAT5 assay, cells were stained using the transcription factor phospho buffer set (BD Pharmingen) according to the manufacturer’s instructions.

The following reagents and antibodies were used: Live/dead fixable dead cell stain kits (blue and violet, Molecular Probes), TruStain FcX anti-mouse CD16/CD32 (93, BioLegend), anti-CD4 PerCP (RM4-5, BD Pharmingen), anti-CD4 BV785 (RM4-5, BioLegend), anti-CD8α V500 and anti-CD8α BUV395 (53-6.7, BD Horizon), anti-CD25 BV421 (7D4, BD Horizon), anti-CD25 FITC (7D4, BD Pharmingen), anti-CD25 PE (PC61.5, eBioscience), anti-CD44 AF700 (IM7, BioLegend), anti-CD49b PE-Cy7 (DX5, eBioscience), anti-CD62L (MEL-14, BD Pharmingen), anti-CD90.2 BV711 (53-2.1, BD OptiBuild), anti CTLA4 PE-Dazzle594 (UC10-4B9, BioLegend), anti-Foxp3 AF488 and anti-Foxp3 AF647 (MF23, BD Pharmingen), anti-ICOS PE (15F9, BioLegend), anti-Ki-67 BV421 (B56, BD Horizon), anti-Ki-67 FITC (B56, BD Pharmingen), anti-KLRG1 BV510 (2F1, BD OptiBuild), anti-Lag3 BV711 (C9B7W, BD Horizon), anti-pSTAT5 PE (47, BD Phosflow), anti-ST2 BV605 (U29-93, BD Optibuild), and anti-TCRβ APC-Cy7 (H57-597, BD Pharmingen). Intravascular labeling in NOD mice was done using anti-CD45.1 PE (A20, eBioscience) administered by retro-orbital injection few minutes before euthanasia (3μg/mouse).

Flow cytometric data were acquired on BD LSRII flow cytometer using BD FACSDIVA software (version 8.0.1) and analyses were performed with FlowJo software (version 10.5.0).

### Histology

Pancreata were harvested immediately after euthanasia, fixed in 10% neutral buffered formalin (VWR), and embedded in paraffin (Leica). 5μm sections (three/mouse, 200μm apart) were stained with hematoxylin and eosin (Leica). Per section, an average of 16 islets was detected from PBS-treated mice vs 29 islets from Fc.Mut24-treated ones. Islets were scored for the severity of insulitis as described using Leica DM2500 microscope. Images were taken with SPOT Insight 4.0 Megapixel CCD digital camera using SPOT software (version 5.1).

### Statistics

Data were graphed and analyzed for statistical significance using GraphPad Prism software (version 8.0.0). For titration curves, EC50 was calculated using nonlinear regression assuming a variable slope (four parameters). For *in vivo* studies, statistical differences among groups were calculated using either a one-way ANOVA or a two-way ANOVA followed by Tukey’s multiple comparisons test. For Fc.IL-2 association with surface CD25 in vitro, statistical differences between Fc.WT and Fc.Mut24 at various time points were determined by repeated measures two-way ANOVA with the Geisser-Greenhouse correction followed by Tukey’s multiple comparisons test. For diabetes studies in NOD mice, statistical differences between treatment groups were calculated using Log-rank (Mantel-Cox) test. Data were presented as the mean ± SD unless otherwise specified and MFI represented median fluorescence intensity. Significance is determined by p ≤ 0.5 (*p<0.05, **p<0.01, ****p<0.0001) and all significant comparisons are shown.

## Supporting information

Supplemental Methods, Figures and Tables for Khoryati et al.

## List of supplementary materials

Fig. S1. Fc.WT shows a comparable activity to mouse recombinant IL-2.

Fig. S2. Fc.mutein activity in Tregs ranges from full potency to complete functional abrogation.

Fig. S3. Fc.muteins bind CD25 with the same affinity as Fc.WT.

Fig. S4. Fc.muteins retain Treg-selectivity across a wide dose range on activated Tcells.

Fig. S5. Fc.Mut24 remains Treg-selective across a wide dose range *in vivo*.

Fig. S6. Fc.Mut24 expands Tregs more specifically and effectively than daily low-dose IL-2 and IL-2ic *in vivo*.

Fig. S7. Sustained Treg enrichment following Fc.Mut24 treatment.

Fig. S8. Reduced Treg selectivity of Fc.Mut24 in pancreatic cells.

Table S1. Panel of Fc.IL-2 muteins.

## Acknowledgments

We thank the Campbell, Long, Buckner and Cerosaletti lab members and C. Pfleger for helpful discussions. We thank C. Stefani and A. Lacy-Hulbert for providing HEK cells and transfection reagents. We thank Benaroya Research Institute core laboratories for technical assistance: P. Johnson and R. Vernon (Histology/Imaging core), K. Arumuganathan, A. Wojno, T. Nguyen (Flow Cytometry core), K. Schwedhelm, J. Thorpe, M. Maerz, and A. Kus (Human Immunophenotyping core), A. Burich, C. Toledano and the vivarium personnel (Animal Resources).

## Funding

This work was funded in part by JDRF 2-SRA-2016-305-S-B and Amgen, Inc. to M.A.G. and NIH grant R01AI136475 to D.J.C.

## Author contributions

L.K., D.J.C., and M.A.G. conceived and designed the study; L.K., M.N.P., M.S., S.K., K.C., J.P., and M.A.G. performed the experiments; L.K., K.C., J.P., M.B., D.J.C., and M.A.G. analyzed and interpreted the data; L.K., D.J.C., and M.A.G. wrote the manuscript. All authors approved the final manuscript.

## Competing interests

M.A.G., K.C. and J.P. own stock in Amgen, Inc.

## Notes

### Summary of Updates

Added new data and expanded introduction and discussion sections of manuscript

## References

1. T. R. Malek, The biology of interleukin-2. Annu. Rev. Immunol. 26, 453–479 (2008).

2. A. K. Abbas, E. Trotta, D. R. Simeonov, A. Marson, J. A. Bluestone, Revisiting IL-2: Biology and therapeutic prospects. Sci. Immunol. 3, eaat1482 (2018).

3. X. Wang, M. Rickert, K. C. Garcia, Structure of the quaternary complex of interleukin-2 with its α, β and γc receptors. Science. 310, 1159–1163 (2005).

4. S. Sakaguchi, N. Sakaguchi, M. Asano, M. Itoh, M. Toda, Immunologic self-tolerance maintained by activated T cells expressing IL-2 receptor alpha-chains (CD25). Breakdown of a single mechanism of self-tolerance causes various autoimmune diseases. J. Immunol. 155, 1151–1164 (1995).

5. J. D. Fontenot, J. P. Rasmussen, M. A. Gavin, A. Y. Rudensky, A function for interleukin 2 in Foxp3-expressing regulatory T cells. Nat Immunol. 6, 1142–1151 (2005).

6. C. Baecher-Allan, J. A. Brown, G. J. Freeman, D. A. Hafler, CD4 + CD25 high Regulatory Cells in Human Peripheral Blood. J. Immunol. 167, 1245–1253 (2001).

7. O. Boyman, J. Sprent, The role of interleukin-2 during homeostasis and activation of the immune system. Nat. Rev. Immunol. 12, 180–190 (2012).

8. D. A. Cantrell, K. A. Smith, Transient expression of interleukin 2 receptors: Consequences for T cell growth. J. Exp. Med. 158, 1895–1911 (1983).

9. H. P. Kim, J. Imbert, W. J. Leonard, Both integrated and differential regulation of components of the IL-2/IL-2 receptor system. Cytokine Growth Factor Rev. 17, 349–366 (2006).

10. A. N. Shatrova, E. V. Mityushova, I. O. Vassilieva, N. D. Aksenov, V. V. Zenin, N. N. Nikolsky, I. I. Marakhova, Time-dependent regulation of IL-2R α-chain (CD25) expression by TCR signal strength and IL-2-induced STAT5 signaling in activated human blood T lymphocytes. PLoS One. 11, e0167215 (2016).

11. M. A. Gavin, J. P. Rasmussen, J. D. Fontenot, V. Vasta, V. C. Manganiello, J. A. Beavo, A. Y. Rudensky, Foxp3-dependent programme of regulatory T-cell differentiation. Nature. 445, 771–775 (2007).

12. Z. Yao, Y. Kanno, M. Kerenyi, G. Stephens, L. Durant, W. T. Watford, A. Laurence, G. W. Robinson, E. M. Shevach, R. Moriggl, L. Hennighausen, C. Wu, J. J. O’Shea, Nonredundant roles for Stat5a/b in directly regulating Foxp3. Blood. 109, 4368–4375 (2007).

13. A. Y. Rudensky, Regulatory T cells and Foxp3. Immunol. Rev. 241, 260–268 (2011).

14. Y. Feng, A. Arvey, T. Chinen, J. Van Der Veeken, G. Gasteiger, A. Y. Rudensky, Control of the inheritance of regulatory T cell identity by a cis element in the Foxp3 locus. Cell. 158, 749–763 (2014).

15. G. C. Sim, L. Radvanyi, The IL-2 cytokine family in cancer immunotherapy. Cytokine Growth Factor Rev. 25, 377–390 (2014).

16. J. M. Wrangle, A. Patterson, C. B. Johnson, D. J. Neitzke, S. Mehrotra, C. E. Denlinger, C. M. Paulos, Z. Li, D. J. Cole, M. P. Rubinstein, IL-2 and beyond in cancer immunotherapy. J. Interf. Cytokine Res. 38, 45–68 (2018).

17. P. Hu, M. Mizokami, G. Ruoff, L. A. Khawli, A. L. Epstein, Generation of low-toxicity interleukin-2 fusion proteins devoid of vasopermeability activity. Blood. 101, 4853–4861 (2003).

18. S. D. Gillies, Y. Lan, T. Hettmann, B. Brunkhorst, Y. Sun, S. O. Mueller, K. M. Lo, A low-toxicity IL-2-based immunocytokine retains antitumor activity despite its high degree of IL-2 receptor selectivity. Clin. Cancer Res. 17, 3673–3685 (2011).

19. J. Laurent, C. Touvrey, S. Gillessen, M. Joffraud, M. Vicari, C. Bertrand, S. Ongarello, B. Liedert, E. Gallerani, J. Beck, A. Omlin, C. Sessa, S. Quaratino, R. Stupp, U. S. Gnad-Vogt, D. E. Speiser, T-cell activation by treatment of cancer patients with EMD 521873 (Selectikine), an IL-2/anti-DNA fusion protein. J. Transl. Med. 11, 5 (2013).

20. A. M. Levin, D. L. Bates, A. M. Ring, C. Krieg, J. T. Lin, L. Su, I. Moraga, M. E. Raeber, G. R. Bowman, P. Novick, V. S. Pande, C. G. Fathman, O. Boyman, K. C. Garcia, Exploiting a natural conformational switch to engineer an interleukin-2 “superkine.” Nature. 484, 529–533 (2012).

21. D. A. Silva, S. Yu, U. Y. Ulge, J. B. Spangler, K. M. Jude, C. Labão-Almeida, L. R. Ali, A. Quijano-Rubio, M. Ruterbusch, I. Leung, T. Biary, S. J. Crowley, E. Marcos, C. D. Walkey, B. D. Weitzner, F. Pardo-Avila, J. Castellanos, L. Carter, L. Stewart, S. R. Riddell, M. Pepper, G. J. L. Bernardes, M. Dougan, K. C. Garcia, D. Baker, De novo design of potent and selective mimics of IL-2 and IL-15. Nature. 565, 186–191 (2019).

22. D. Klatzmann, A. K. Abbas, The promise of low-dose interleukin-2 therapy for autoimmune and inflammatory diseases. Nat. Rev. Immunol. 15, 283–294 (2015).

23. C. Ye, D. Brand, S. G. Zheng, Targeting IL-2: an unexpected effect in treating immunological diseases. Signal Transduct. Target. Ther. 3, 2 (2018).

24. E. Seelig, J. Howlett, L. Porter, L. Truman, J. Heywood, J. Kennet, E. L. Arbon, K. Anselmiova, N. M. Walker, R. Atkar, M. L. Pekalski, E. Rytina, M. Evans, L. S. Wicker, J. A. Todd, A. P. Mander, S. Bond, F. Waldron-Lynch, The DILfrequency study is an adaptive trial to identify optimal IL-2 dosing in patients with type 1 diabetes. JCI Insight. 3, e99306 (2018).

25. L. B. Peterson, C. J. M. Bell, S. K. Howlett, M. L. Pekalski, K. Brady, H. Hinton, D. Sauter, J. A. Todd, P. Umana, O. Ast, I. Waldhauer, A. Freimoser-Grundschober, E. Moessner, C. Klein, R. J. Hosse, L. S. Wicker, A long-lived IL-2 mutein that selectively activates and expands regulatory T cells as a therapy for autoimmune disease. J. Autoimmun. 95, 1–14 (2018).

26. A. B. Shanafelt, Y. Lin, M. C. Shanafelt, C. P. Forte, N. Dubois-Stringfellow, C. Carter, J. A. Gibbons, S. L. Cheng, K. A. Delaria, R. Fleischer, J. M. Greve, R. Gundel, K. Harris, R. Kelly, B. Koh, Y. Li, L. Lantz, P. Mak, L. Neyer, M. J. Plym, S. Roczniak, D. Serban, J. Thrift, L. Tsuchiyama, M. Wetzel, M. Wong, A. Zolotorev, A T-cell-selective interleukin 2 mutein exhibits potent antitumor activity and is well tolerated in vivo. Nat. Biotechnol. 18, 1197–1202 (2000).

27. S. M. Zurawski, G. Zurawski, Mouse interleukin-2 structure-function studies: substitutions in the first alpha-helix can specifically inactivate p70 receptor binding and mutations in the fifth alpha-helix can specifically inactivate p55 receptor binding. EMBO J. 8, 2583–2590 (1989).

28. S. M. Zurawski, J. L. Imler, G. Zurawski, Partial agonist/antagonist mouse interleukin-2 proteins indicate that a third component of the receptor complex functions in signal transduction. EMBO J. 9, 3899–3905 (1990).

29. S. M. Zurawski, F. Vega, Jr., E. L. Doyle, B. Huyghe, K. Flaherty, D. B. McKay, G. Zurawski, Definition and spatial location of mouse interleukin-2 residues that interact with its heterotrimeric receptor. EMBO J. 12, 5113–5119 (1993).

30. F. W. Jacobsen, R. Stevenson, C. Li, H. Salimi-Moosavi, L. Liu, J. Wen, Q. Luo, K. Daris, L. Buck, S. Miller, S. Y. Ho, W. Wang, Q. Chen, K. Walker, J. Wypych, L. Narhi, K. Gunasekaran, Engineering an IgG scaffold lacking effector function with optimized developability. J. Biol. Chem. 292, 1865–1875 (2017).

31. S. C. Wuest, J. H. Edwan, J. F. Martin, S. Han, J. S. A. Perry, C. M. Cartagena, E. Matsuura, D. Maric, T. A. Waldmann, B. Bielekova, A role for interleukin-2 trans-presentation in dendritic cell-mediated T cell activation in humans, as revealed by daclizumab therapy. Nat. Med. 17, 604–609 (2011).

32. Q. Tang, J. Y. Adams, C. Penaranda, K. Melli, E. Piaggio, E. Sgouroudis, C. A. Piccirillo, B. L. Salomon, J. A. Bluestone, Central Role of Defective Interleukin-2 Production in the Triggering of Islet Autoimmune Destruction. Immunity. 28, 687–697 (2008).

33. C. J. M. Bell, Y. Sun, U. M. Nowak, J. Clark, S. Howlett, M. L. Pekalski, X. Yang, O. Ast, I. Waldhauer, A. Freimoser-Grundschober, E. Moessner, P. Umana, C. Klein, R. J. Hosse, L. S. Wicker, L. B. Peterson, Sustained in vivo signaling by long-lived IL-2 induces prolonged increases of regulatory T cells. J. Autoimmun. 56, 66–80 (2015).

34. J. A. Pearson, F. S. Wong, L. Wen, The importance of the Non Obese Diabetic (NOD) mouse model in autoimmune diabetes. J. Autoimmun. 66, 76–88 (2016).

35. C. Baecher-Allan, D. A. Hafler, Human regulatory T cells and their role in autoimmune disease. Immunol.Rev. 212, 203–216 (2006).

36. T. R. Malek, A. Yu, L. Zhu, T. Matsutani, D. Adeegbe, A. L. Bayer, IL-2 family of cytokines in T regulatory cell development and homeostasis. J. Clin. Immunol. 28, 635–639 (2008).

37. D. Saadoun, M. Rosenzwajg, F. Joly, A. Six, F. Carrat, V. Thibault, D. Sene, P. Cacoub, D. Klatzmann, Regulatory T-cell responses to low-dose interleukin-2 in HCV-induced vasculitis. N. Engl. J. Med. 365, 2067–2077 (2011).

38. J. Koreth, K. I. Matsuoka, H. T. Kim, S. M. McDonough, B. Bindra, E. P. Alyea, III, P. Armand, C. Cutler, V. T. Ho, N. S. Treister, D. C. Bienfang, S. Prasad, D. Tzachanis, R. M. Joyce, D. E. Avigan, J. H. Antin, J. Ritz, R. J. Soiffer, Interleukin-2 and regulatory T cells in graft-versus-host disease. N. Engl. J. Med. 365, 2055–2066 (2011).

39. A. A. Kennedy-Nasser, S. Ku, P. Castillo-Caro, Y. Hazrat, M.-F. Wu, H. Liu, J. Melenhorst, A. J. Barrett, S. Ito, A. Foster, B. Savoldo, E. Yvon, G. Carrum, C. A. Ramos, R. A. Krance, K. Leung, H. E. Heslop, M. K. Brenner, C. M. Bollard, Ultra low-dose IL-2 for GVHD prophylaxis after allogeneic hematopoietic stem cell transplantation mediates expansion of regulatory T cells without diminishing antiviral and antileukemic activity. Clin. Cancer Res. 20, 2215–2225 (2014).

40. J. S. Whangbo, H. T. Kim, N. Mirkovic, L. Leonard, S. Poryanda, S. Silverstein, S. Kim, C. G. Reynolds, S. C. Rai, K. Verrill, M. A. Lee, S. Margossian, C. Duncan, L. Lehmann, J. Huang, S. Nikiforow, E. P. Alyea, P. Armand, C. S. Cutler, V. T. Ho, B. R. Blazar, J. H. Antin, R. J. Soiffer, J. Ritz, J. Koreth, Dose-escalated interleukin-2 therapy for refractory chronic graft-versus-host disease in adults and children. Blood Adv. 3, 2550–2561 (2019).

41. E. Castela, F. Le Duff, C. Butori, M. Ticchioni, P. Hofman, P. Bahadoran, J.-P. Lacour, T. Passeron, Effects of low-dose recombinant interleukin 2 to promote T-regulatory cells in alopecia areata. JAMA dermatology. 150, 751 (2014).

42. J. Y. Humrich, C. von Spee-Mayer, E. Siegert, T. Alexander, F. Hiepe, A. Radbruch, G.-R. Burmester, G. Riemekasten, Rapid induction of clinical remission by low-dose interleukin-2 in a patient with refractory SLE. Ann. Rheum. Dis. 74, 792 (2015).

43. C. Von Spee-Mayer, E. Siegert, D. Abdirama, A. Rose, A. Klaus, T. Alexander, P. Enghard, B. Sawitzki, F. Hiepe, A. Radbruch, G. R. Burmester, G. Riemekasten, J. Y. Humrich, Low-dose interleukin-2 selectively corrects regulatory T cell defects in patients with systemic lupus erythematosus. Ann. Rheum. Dis. 75, 1407–1415 (2016).

44. J. He, X. Zhang, Y. Wei, X. Sun, Y. Chen, J. Deng, Y. Jin, Y. Gan, X. Hu, R. Jia, C. Xu, Z. Hou, Y. A. Leong, L. Zhu, J. Feng, Y. An, Y. Jia, C. Li, X. Liu, H. Ye, L. Ren, R. Li, H. Yao, Y. Li, S. Chen, X. Zhang, Y. Su, J. Guo, N. Shen, E. F. Morand, D. Yu, Z. Li, Low-dose interleukin-2 treatment selectively modulates CD4+ T cell subsets in patients with systemic lupus erythematosus. Nat. Med. 22, 991–993 (2016).

45. J. Y. Humrich, C. von Spee-Mayer, E. Siegert, M. Bertolo, A. Rose, D. Abdirama, P. Enghard, B. Stuhlmüller, B. Sawitzki, D. Huscher, F. Hiepe, T. Alexander, E. Feist, A. Radbruch, G. R. Burmester, G. Riemekasten, Low-dose interleukin-2 therapy in refractory systemic lupus erythematosus: an investigator-initiated, single-centre phase 1 and 2a clinical trial. Lancet Rheumatol. 1, e44–e54 (2019).

46. J. He, R. Zhang, M. Shao, X. Zhao, M. Miao, J. Chen, J. Liu, X. Zhang, X. Zhang, Y. Jin, Y. Wang, S. Zhang, L. Zhu, A. Jacob, R. Jia, X. You, X. Li, C. Li, Y. Zhou, Y. Yang, H. Ye, Y. Liu, Y. Su, N. Shen, J. Alexander, J. Guo, J. Ambrus, X. Lin, D. Yu, X. Sun, Z. Li, Efficacy and safety of low-dose IL-2 in the treatment of systemic lupus erythematosus: a randomised, double-blind, placebo-controlled trial. Ann. Rheum. Dis. 79, 141–149 (2020).

47. M. Rosenzwajg, R. Lorenzon, P. Cacoub, H. P. Pham, F. Pitoiset, K. El Soufi, C. Rlbet, C. Bernard, S. Aractingi, B. Banneville, L. Beaugerie, F. Berenbaum, J. Champey, O. Chazouilleres, C. Corpechot, B. Fautrel, A. Mekinian, E. Regnier, D. Saadoun, J. E. Salem, J. Sellam, P. Seksik, A. Daguenel-Nguyen, V. Doppler, J. Mariau, E. Vicaut, D. Klatzmann, Immunological and clinical effects of low-dose interleukin-2 across 11 autoimmune diseases in a single, open clinical trial. Ann. Rheum. Dis. 78, 209–217 (2019).

48. J. Koreth, J. Ritz, G. C. Tsokos, A. Pugliese, T. R. Malek, M. Rosenzwajg, D. Klatzmann, Low-dose Interleukin-2 in the Treatment of Autoimmune Disease. Oncol. Hematol. Rev. 10, 157–163 (2014).

49. S. A. Long, K. Cerosaletti, P. L. Bollyky, M. Tatum, H. Shilling, S. Zhang, Z. Zhang, C. Pihoker, S. Sanda, C. Greenbaum, J. H. Buckner, Defects in IL-2R Signaling Contribute to Diminished Regulatory T-Cells of Type 1 Diabetic Subjects. Diabetes. 59, 407–415 (2010).

50. D. J. Stauber, E. W. Debler, P. A. Horton, K. A. Smith, I. A. Wilson, Crystal structure of the IL-2 signaling complex: Paradigm for a heterotrimeric cytokine receptor. Proc. Natl. Acad. Sci. U. S. A. 103, 2788–2793 (2006).

51. N. Tchao, K. S. Gorski, T. Yuraszeck, S. J. Sohn, K. Ishida, H. Wong, K. Park, Amg 592 is an investigational IL-2 mutein that induces highly selective expansion of regulatory T cells. Blood. 130, 696 (2017).

52. K. S. Gorski, J. Stern, Y.-H. Hsu, A. Anderson, M. Boedigheimer, N. Tchao, Phenotype of foxp3+ regulatory t-cells expanded by the il-2 mutein, amg 592 in healthy subjects in phase 1, first-in-human study. Ann. Rheum. Dis. 77, 243 (2018).

53. K. C. Herold, B. N. Bundy, S. A. Long, J. A. Bluestone, L. A. DiMeglio, M. J. Dufort, S. E. Gitelman, P. A. Gottlieb, J. P. Krischer, P. S. Linsley, J. B. Marks, W. Moore, A. Moran, H. Rodriguez, W. E. Russell, D. Schatz, J. S. Skyler, E. Tsalikian, D. K. Wherrett, A.-G. Ziegler, C. J. Greenbaum, An anti-CD3 antibody, teplizumab, in relatives at risk for type 1 diabetes. N. Engl. J. Med. 381, 603–613 (2019).

