## Supplemental Methods, Figures and Tables for Khoryati et al. for "Regulatory T cell expansion by a highly CD25-dependent IL-2 mutein arrests ongoing autoimmunity"

**This PDF file includes:**

Figs. S1 to S8

Table S1

**Supplementary Figures**

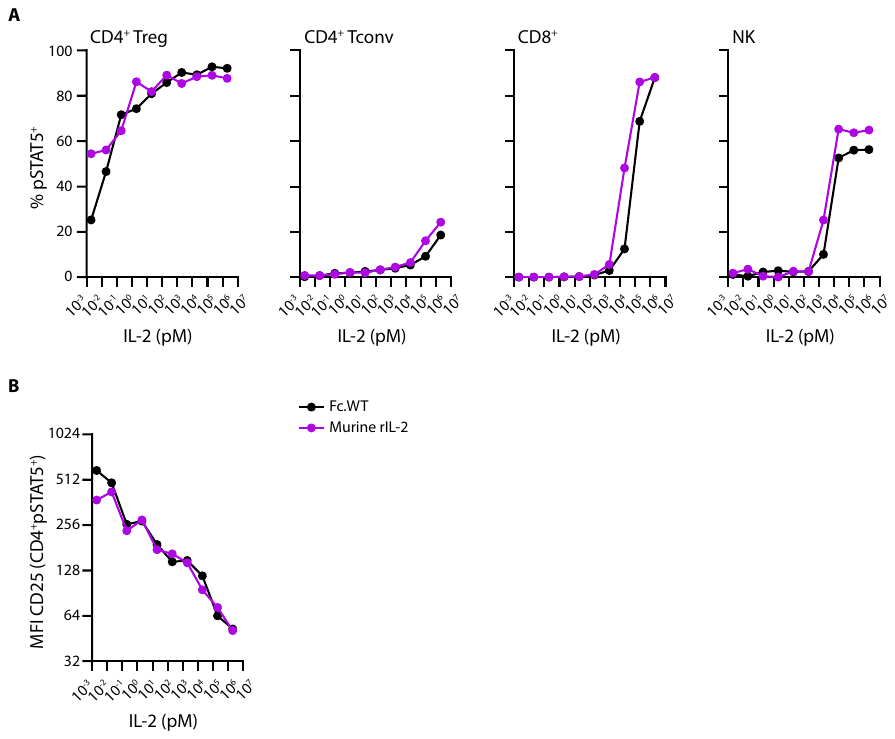

**Fig. S1. Fc.WT shows a comparable activity to mouse recombinant IL-2.** B6 Splenocytes were stimulated with either Fc.WT or mouse recombinant IL-2 (**A**) pSTAT5 dose-response curves for the indicated populations. (**B**) CD25 median fluorescence intensity on CD4^+^pSTAT5^+^ cells.

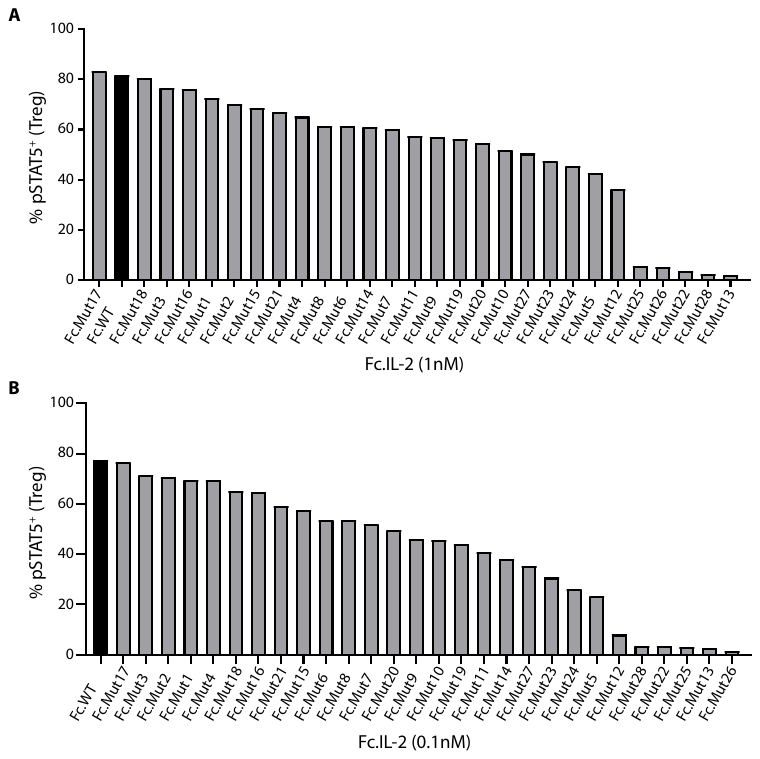

**Fig. S2. Fc.mutein activity in Tregs ranges from full potency to complete functional abrogation.** B6 Splenocytes were stimulated with transfection supernatants for the indicated Fc.IL-2 fusion proteins. Frequencies of pSTAT5^+^ cells in Tregs at **(A)** 1nM and **(B)** 0.1nM of Fc.IL-2 are plotted.

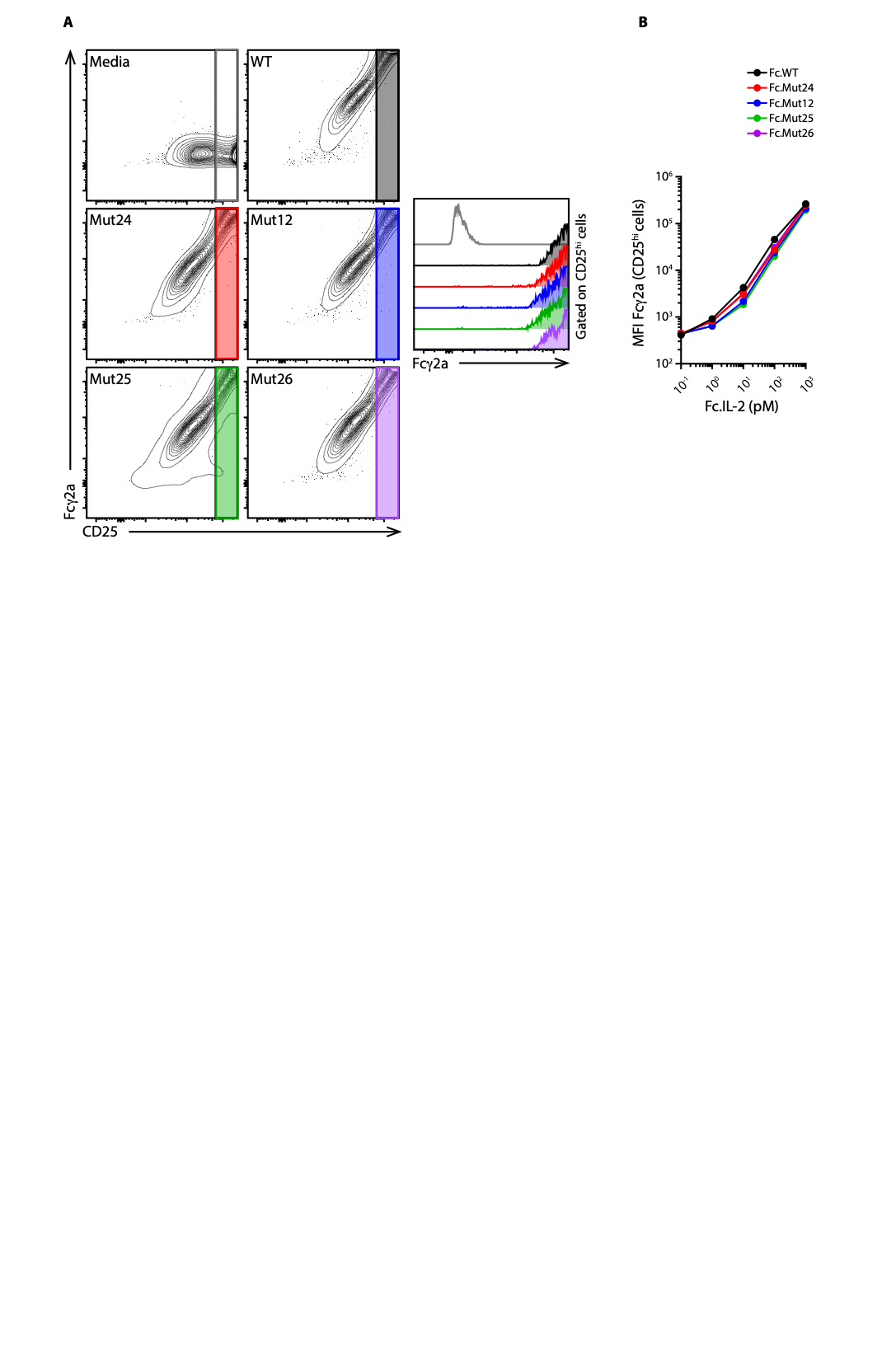

**Fig. S3. Fc.muteins bind CD25 with the same affinity as Fc.WT.** CD25-expressing HEK 293T/17 cells were coated with titrated purified Fc.IL-2 proteins. **(A)** Left: Representative staining for surface Fc.IL-2 and CD25 on CD25-expressing HEK cells. Right: Analysis of Fcγ2a staining on CD25^hi^ cells (coating with 1nM Fc.IL-2). (**B**) Analyses of median fluorescence intensity of Fcγ2a on CD25^hi^ cells.

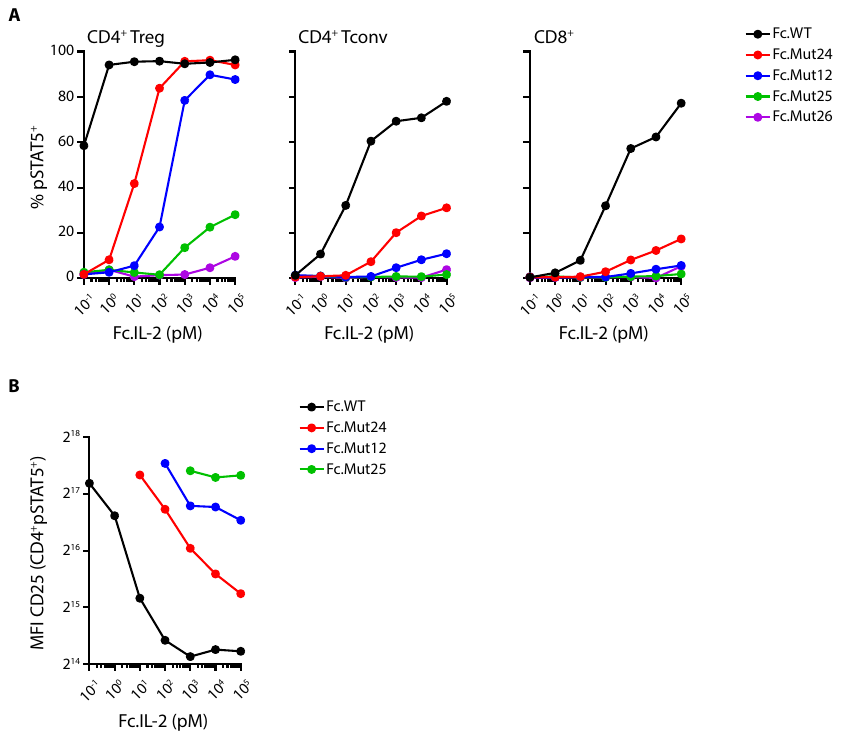

**Fig. S4. Fc.muteins retain Treg-selectivity across a wide dose range on activated Tcells.** Anti-CD3 activated B6 splenocytes were stimulated with the indicated purified Fc.IL-2 fusion proteins. (**A)** pSTAT5 dose-response curves for the indicated populations. (**B**) CD25 median fluorescence intensity on CD4^+^pSTAT5^+^ cells.

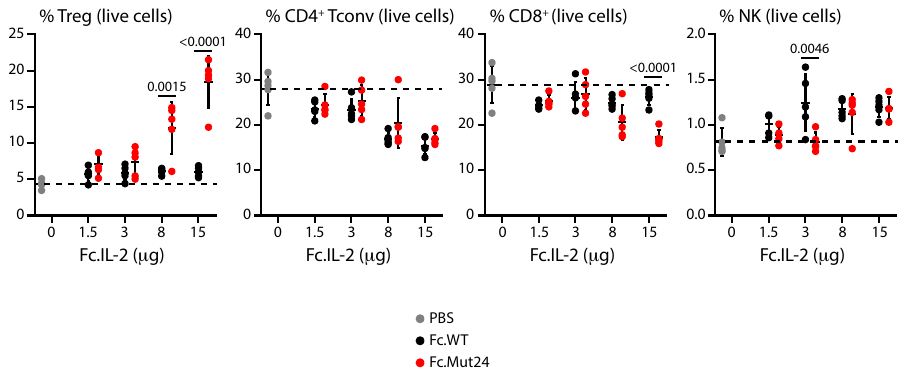

**Fig. S5.** **Fc.Mut24 remains Treg-selective across a wide dose range *in vivo*.** B6 Mice were treated on day 0 with either PBS or Fc.IL-2 and inguinal lymph nodes were harvested four days later for analysis by flow cytometry. Frequencies of the indicated populations in live cells are plotted. Data shown as mean ± SD, significance determined by two-way ANOVA followed by Tukey post-test. Representative of at least two independent experiments with n=5 mice/group.

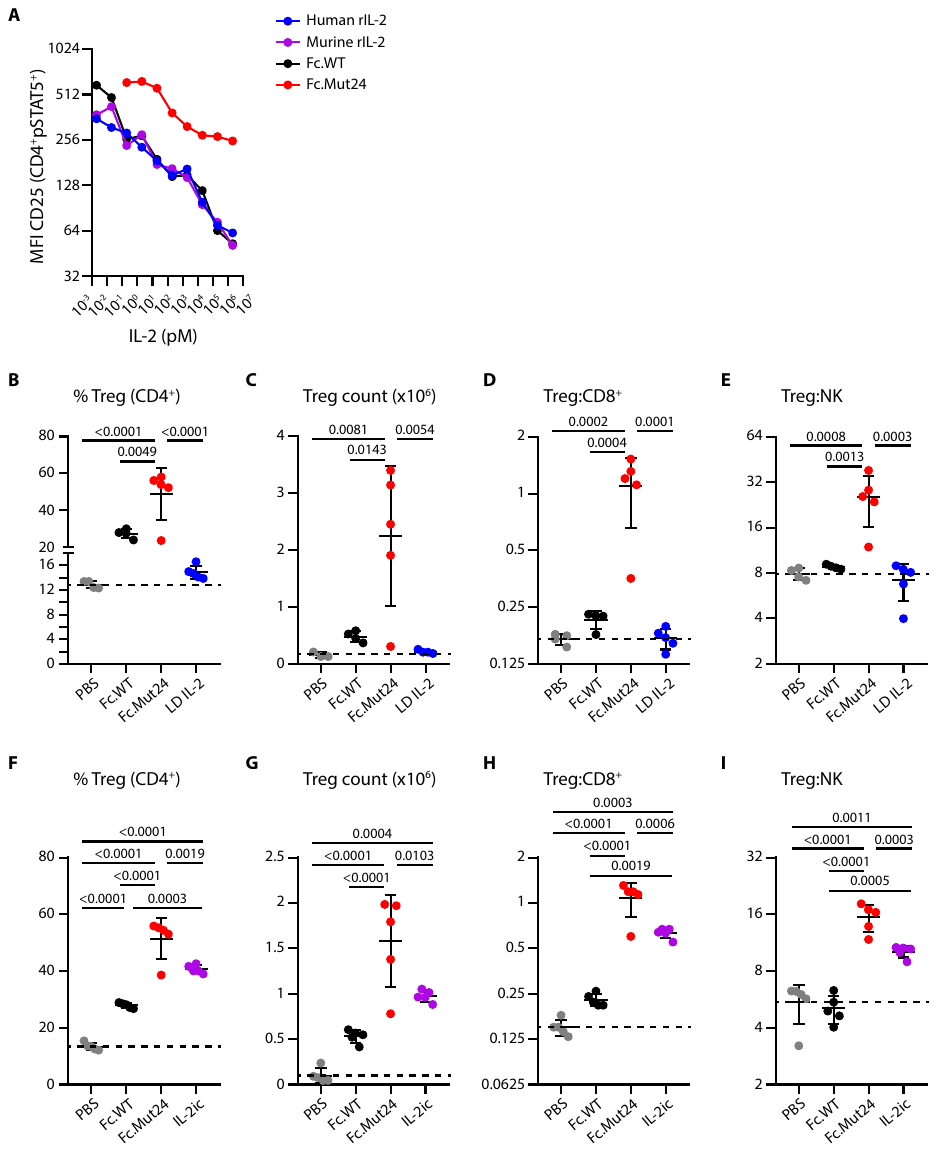
 **Fig. S6.** **Fc.Mut24 expands Tregs more specifically and effectively than daily low-dose IL-2 and IL-2ic *in vivo*.** (A) B6 Splenocytes were stimulated with human recombinant IL-2, mouse recombinant IL-2, or Fc.IL-2. CD25 median fluorescence intensity on CD4^+^pSTAT5^+^ cells is shown. (B to E) B6 Mice were treated on day 0 with either a single injection of Fc.IL-2 (15μg) or four consecutive daily injections of low-dose human recombinant IL-2 (25,000 IU/injection, day 0 through day 3). Control mice were administered PBS. On day 4 after initiation of the treatment, inguinal lymph nodes (iLN) were harvested for analysis by flow cytometry. (**B**) Treg frequencies, (C) Treg count (from 2 pooled iLN), and ratios of (**D**) Treg:CD8^+^ and (**E**) Treg:NK are summarized. (G to I) B6 Mice were treated on day 0 with a single dose of either Fc.IL-2 (15μg) or equimolar amount of mouse recombinant IL-2 complexed with JES6-1A12 antibody (IL-2ic). Control mice received PBS. Four days later, iLN were harvested for analysis. (**F**) Treg frequencies, (G) Treg count (from 2 pooled iLN), and ratios of (**H**) Treg:CD8^+^ and (**I**) Treg:NK are summarized. Data shown as mean ± SD with n=5 mice/group. Significance determined by one-way ANOVA followed by Tukey post-test.

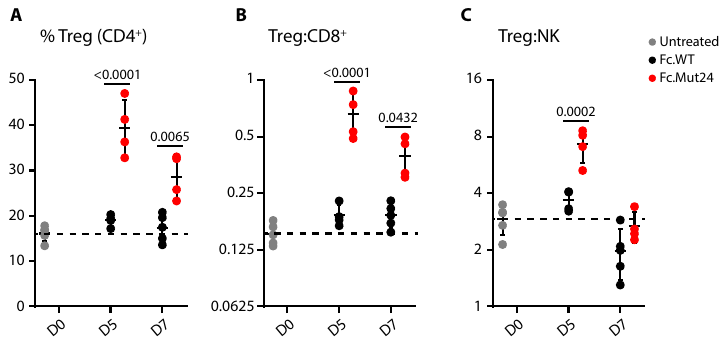

**Fig. S7. Sustained Treg enrichment following Fc.Mut24 treatment.** B6 Mice were treated with Fc.WT or Fc.Mut24 (15μg) on day 0 and iLN were harvested on day 5 and 7 for analysis. (**A**) Treg frequencies and ratios of (**C**) Treg:CD8^+^ and (**D**) Treg:NK are plotted. Data shown as mean ± SD, significance determined by two-way ANOVA followed by Tukey post-test. Representative of two independent experiments with n=4-5 mice/group.

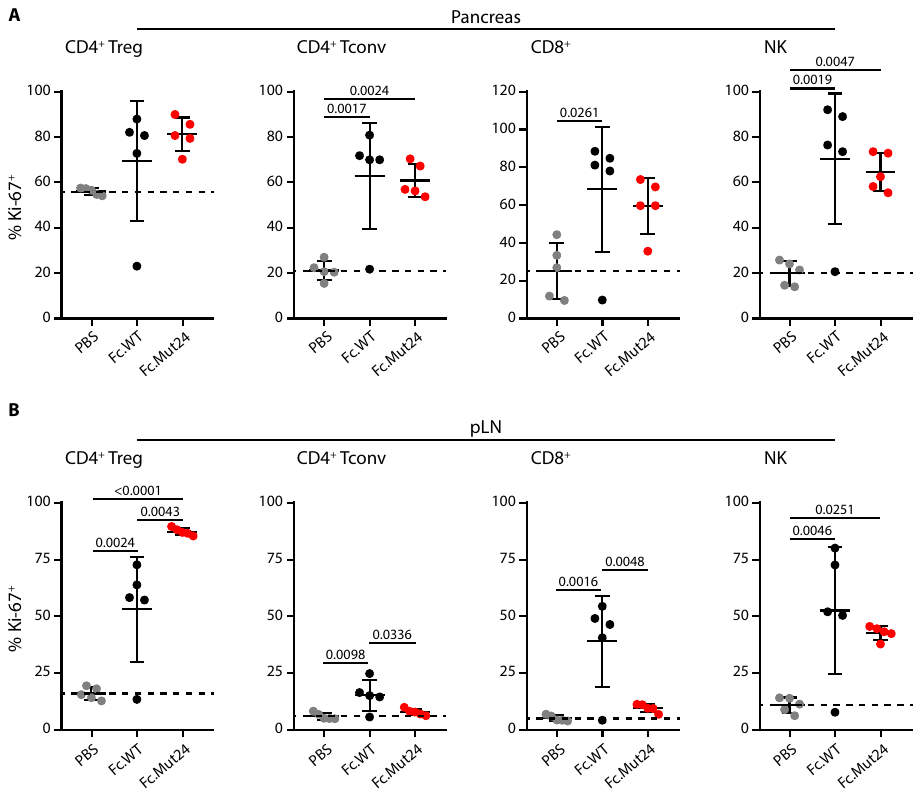

**Fig. S8. Reduced Treg selectivity of Fc.Mut24 in pancreatic cells.** Female NOD mice were treated on day 0 with PBS, Fc.WT or Fc.Mut24 (15μg) and tissues were harvested four days later. Percentages of Ki-67^+^ cells in the indicated populations in the **(A)** pancreas and **(B)** pancreatic lymph node are plotted. Data shown as mean ± SD. Representative of at least two independent experiments with n=5 mice/group.

| **Mutein** | **Mutation (s)** |
| --- | --- |
| **Fc.Mut1** | **Q30K** |
| **Fc.Mut2** | **Q30R** |
| **Fc.Mut3** | **Q30D** |
| **Fc.Mut4** | **Q30E** |
| **Fc.Mut5** | **D34R** |
| **Fc.Mut6** | **D34G** |
| **Fc.Mut7** | **D34T** |
| **Fc.Mut8** | **D34N** |
| **Fc.Mut9** | **D34Q** |
| **Fc.Mut10** | **D34H** |
| **Fc.Mut11** | **N103R** |
| **Fc.Mut12** | **N103K** |
| **Fc.Mut13** | **N103E** |
| **Fc.Mut14** | **N103H** |
| **Fc.Mut15** | **V106D** |
| **Fc.Mut16** | **V106E** |
| **Fc.Mut17** | **V106G** |
| **Fc.Mut18** | **V106K** |
| **Fc.Mut19** | **M33D-V106K** |
| **Fc.Mut20** | **M33E-V106K** |
| **Fc.Mut21** | **M33G-V106K** |
| **Fc.Mut22** | **D34K-N103K-V106A** |
| **Fc.Mut23** | **D34K** |
| **Fc.Mut24** | **N103R-V106D** |
| **Fc.Mut25** | **D34Q-V106D** |
| **Fc.Mut26** | **D34H-V106D** |
| **Fc.Mut27** | **M33D-V106D** |
| **Fc.Mut28** | **D34K-N103E-V106K** |

**Table S1. Panel of Fc.IL-2 muteins.**
